# Mutual antagonism between glucocorticoid and canonical Wnt signaling pathways in B-cell acute lymphoblastic leukemia

**DOI:** 10.1101/2023.01.20.524798

**Authors:** Brennan P. Bergeron, Kelly R. Barnett, Kashi Raj Bhattarai, Robert J. Mobley, Baranda S. Hansen, Anthony Brown, Kiran Kodali, Anthony A. High, Sima Jeha, Ching-Hon Pui, Junmin Peng, Shondra M. Pruett-Miller, Daniel Savic

## Abstract

Glucocorticoids (GCs; i.e., steroids) are important chemotherapeutic agents in the treatment of B-cell precursor acute lymphoblastic leukemia (B-ALL) and *de novo* GC resistance predicts relapse and poor clinical outcome in patients. Glucocorticoids induce B-ALL cell apoptosis through activation of glucocorticoid receptor (GR), a ligand-induced nuclear receptor transcription factor (TF). We previously identified disruptions to glucocorticoid receptor (GR)-bound *cis*-regulatory elements controlling *TLE1* expression in GC-resistant primary B-ALL cells from patients. *TLE1* is a GC-response gene up-regulated by steroids and functions as a canonical Wnt signaling repressor. To better understand the mechanistic relationship between GC signaling and canonical Wnt signaling, we performed diverse functional analyses that identified extensive crosstalk and mutual antagonism between these two signaling pathways in B-ALL. We determined that crosstalk and antagonism was driven by the binding of GR and the canonical Wnt signaling TFs LEF1 and TCF7L2 to overlapping sets of *cis*-regulatory elements associated with genes impacting cell death and cell proliferation, and was further accompanied by overlapping and opposing transcriptional programs. Our data additionally suggest that *cis*-regulatory disruptions at *TLE1* are linked to GC resistance through a dampening of the GC response and GC-mediated apoptosis via enhanced canonical Wnt signaling. As a result of the extensive genomic and gene regulatory connectivity between these two signaling pathways, our data supports the importance of canonical Wnt signaling in mediating GC resistance in B-ALL.

## To the Editor

Glucocorticoids (GCs; i.e., steroids) are important chemotherapeutic agents in the treatment of B-cell precursor acute lymphoblastic leukemia (B-ALL)^1^ and *de novo* GC resistance predicts relapse and poor clinical outcome^2,3^. Glucocorticoids induce B-ALL cell apoptosis through activation of glucocorticoid receptor (GR), a ligand-induced nuclear receptor transcription factor^4,5^. We previously identified disruptions to glucocorticoid receptor (GR)-bound *cis*-regulatory elements controlling *TLE1* expression, a GC-response gene up-regulated by steroids, as a novel mechanism impacting GC resistance in primary B-ALL cells from patients^6^. Supporting our findings, *TLE1* expression is associated with GC resistance in primary B-ALL cells from patients, with 40% of GC-resistant patient samples harboring low *TLE1* expression that further correlates with treatment response (**Supplemental Figure 1**)^7,8^. In this regard, *TLE1* was also identified as an indicator of adverse prognosis in patients with ALL^9^. Because TLE1 functions as a repressor of canonical Wnt signaling^10^, and because investigations in other cell systems have suggested interaction between GC and canonical Wnt signaling pathways^11-13^, we investigated mechanisms connecting TLE1 to GC resistance and the extent of crosstalk between GC and canonical Wnt signaling pathways in B-ALL.

We first examined the functional effects of a homozygous knockout (KO) of TLE1 on GC and canonical Wnt signaling using CRISPR-Cas9 genome editing in the Nalm6 human B-ALL cell line (**Figures 1A-B**). Consistent with our previous observations^6^, we found TLE1 KO led to significantly increased resistance to GCs (i.e., prednisolone), with >5-fold increase in prednisolone LC_50_ compared to wild-type (WT) cells (**Figure 1C**). To better understand the broader effects of TLE1 disruption on GC signaling, we conducted RNA-seq in WT and TLE1 KO Nalm6 cells treated with prednisolone or vehicle control for 24 hours. Differentially expressed gene (DEG) analyses between WT and TLE1 KO cells in the presence or absence of prednisolone identified an enrichment for apoptotic signaling and cell cycle pathways at TLE1 KO repressed genes, and diverse metabolic pathways for TLE1 KO activated genes (**Figure 1D** and **supplemental Figure 2** and **supplemental Table 1**). Similar results were obtained using dexamethasone (**Figure 1E**). To identify *TLE1*-dependent changes to canonical Wnt signaling, both WT and *TLE1* KO cells were transfected with the M50 Super 8xTOPFlash reporter plasmid^14^ and treated for 24 hours with a canonical Wnt signaling agonist (CHIR-99021) or vehicle control. This assay measures beta-catenin-driven luciferase activity as an indicator of endogenous canonical Wnt signaling. Under both treatment conditions, *TLE1* KO cells showed increased canonical Wnt activity compared to WT cells (**Figure 1F**), consistent with the role of TLE1 as a repressor of canonical Wnt signaling^10^.

**Figure 1.**
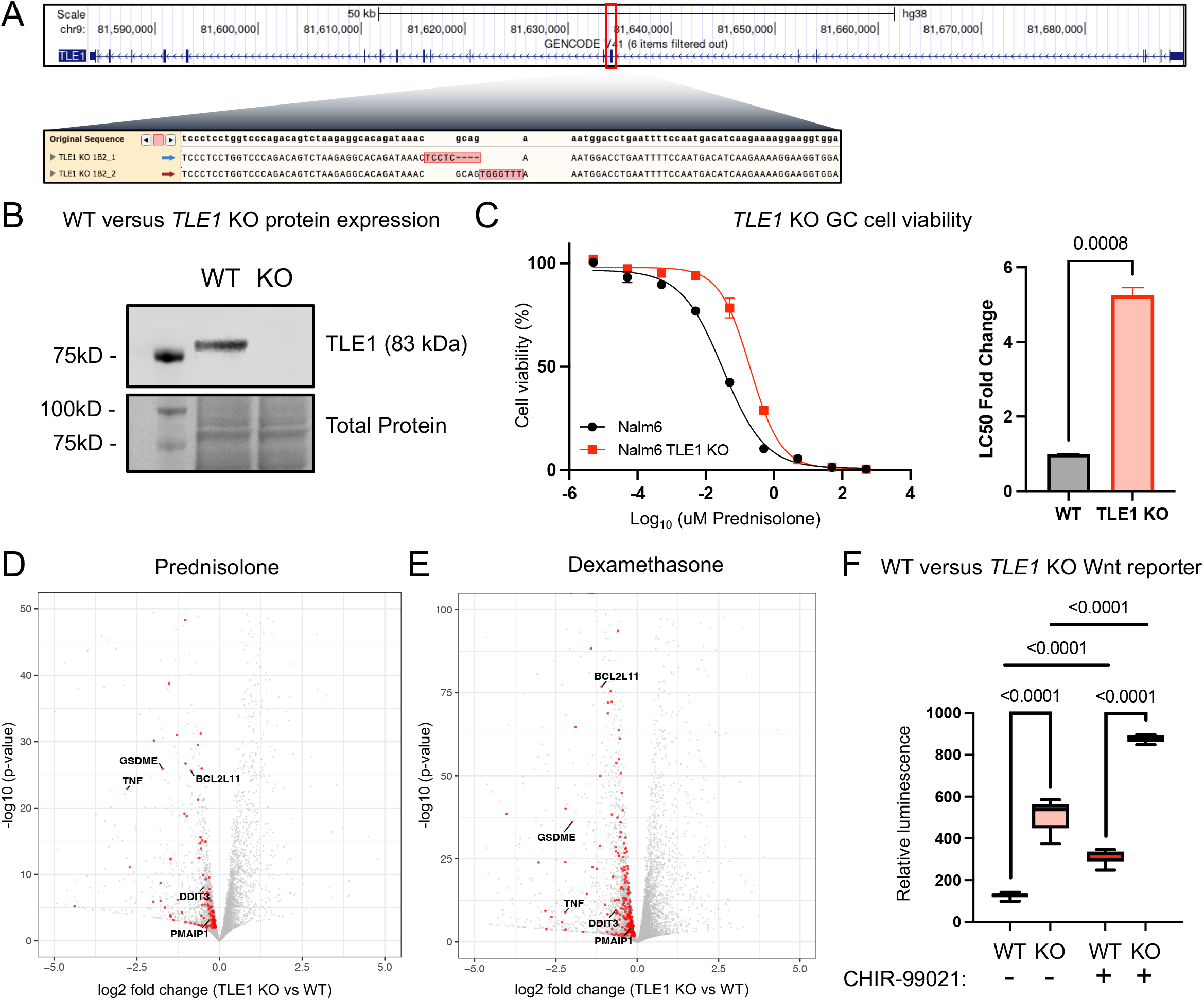
**(A)** Schematic of CRISPR/Cas9 genome editing to exon 7 of the *TLE1* gene with deep sequencing confirmation of edit to both strands in Nalm6 cells **(B)** Western blot showing protein expression of TLE1 (∼83kD; Abcam ab183742) in both TLE1 knockout Nalm6 (KO) and Nalm6 cells (WT). Ponceau staining for total protein below. **(C)** Prednisolone drug response curves in TLE1 knockout Nalm6 cells (red) and wild-type Nalm6 cells (black) after 72-hours of prednisolone treatment is shown at the left, n = 3 per group. Concentrations of prednisolone used were: 5pM, 50pM, 0.5nM, 5nM, 50nM, 0.5uM, 5uM, 50uM, and 500uM. LC50 fold change in *TLE1* gene knockout (TLE1 KO) Nalm6 cells compared to wild-type (WT) Nalm6 cells is shown at the right. **(D-E)** Volcano plots showing differentially expressed genes between WT and TLE1 KO cells after 24 hours of prednisolone (5uM) treatment **(D)** or dexamethasone (100nM) treatment **(E)**. Genes involved in apoptotic pathways are shown in red and several notable genes are highlighted. **(F)** Beta-catenin luciferase reporter assay for wild-type Nalm6 (WT) and *TLE1* KO (KO) cells treated with or without CHIR-99021 (0.5uM) for 24-hours, n = 6 per group.

We next assessed GC and canonical Wnt signaling pathway crosstalk in B-ALL cells independent of *TLE1* KO. Two human B-ALL cell lines (Nalm6 and 697) were treated with different prednisolone concentrations for 72 hours in the presence or absence of Wnt agonist CHIR-99021. At all GC concentrations and in both cell lines, co-treatment with the Wnt agonist significantly increased cellular viability relative to prednisolone-only treatment (**supplemental Figure 3)**. Similar results were also obtained using dexamethasone (**supplemental Figure 4**). We additionally confirmed this effect through *ex vivo* GC drug viability studies in B-ALL patient-derived xenograft cell samples (**supplemental Figure 5)**. These findings suggest that Wnt activation opposes GC-induced apoptosis, which is consistent with the role for canonical Wnt signaling in cell proliferation and cancer predisposition^15^, and prior findings from B-ALL cell viability experiments using Wnt antagonist^16^. Concordant with these data, we identified synergy from co-treatment with prednisolone or dexamethasone and canonical Wnt antagonists (**supplemental Figure 6**), and in TLE1 KO GC-resistant cells (**supplemental Figure 7**)^17^. To further determine the extent that GCs impact canonical Wnt signaling, we measured beta-catenin-driven luciferase reporter activity in B-ALL cells treated with prednisolone and/or CHIR-99021. We found that prednisolone treatment significantly dampened canonical Wnt signaling in the presence or absence of Wnt agonist, with consistent results also observed in TLE1 *KO* cells (**supplemental Figure 8**). Crosstalk between GC and canonical Wnt signaling was also observed in T-cell ALL (T-ALL; **supplemental Figures 9-11**). Collectively, these data indicate crosstalk and antagonism between these two signaling pathways.

We further investigated genomic mechanisms that promote antagonism between GC and canonical Wnt signaling pathways. Using Nalm6 prednisolone- and dexamethasone-response genes, we find that in addition to up-regulation of the *TLE1* canonical Wnt repressor gene, both GCs repressed the canonical Wnt activating transcription factor (TF) gene *LEF1* (**supplemental Figure 12**). Consistent effects on *TLE1* and *LEF1* expression were also reported in a panel of 19 human B-ALL cell lines after dexamethasone treatment^18^. We therefore asked if a 24-hour GC treatment resulted in decreased genome occupancy of LEF1. Using the CUT&RUN assay, we determined that LEF1 binding was decreased at 53% (32105) and increased at only 1.2% (730) of sites after 24 hours of prednisolone treatment (**Figures 2A-B**). Strikingly, we also uncovered that 58% of GR binding sites [previously mapped in ^6^] overlap with LEF1 binding sites (examples in **Figure 2C**), and genes associated with these overlapping sites were significantly enriched for cell proliferation and apoptotic signaling pathways. Consistent results were also observed for the canonical Wnt activating TF TCF7L2 using CUT&RUN, with 91% of differentially bound TCF7L2 sites exhibiting decreased occupancy after prednisolone treatment and 20% of GR binding events overlapping TCF7L2 binding sites at genes enriched for apoptotic signaling (FDR=1.2−10^−8^; **supplemental Figure 13**).

**Figure 2.**
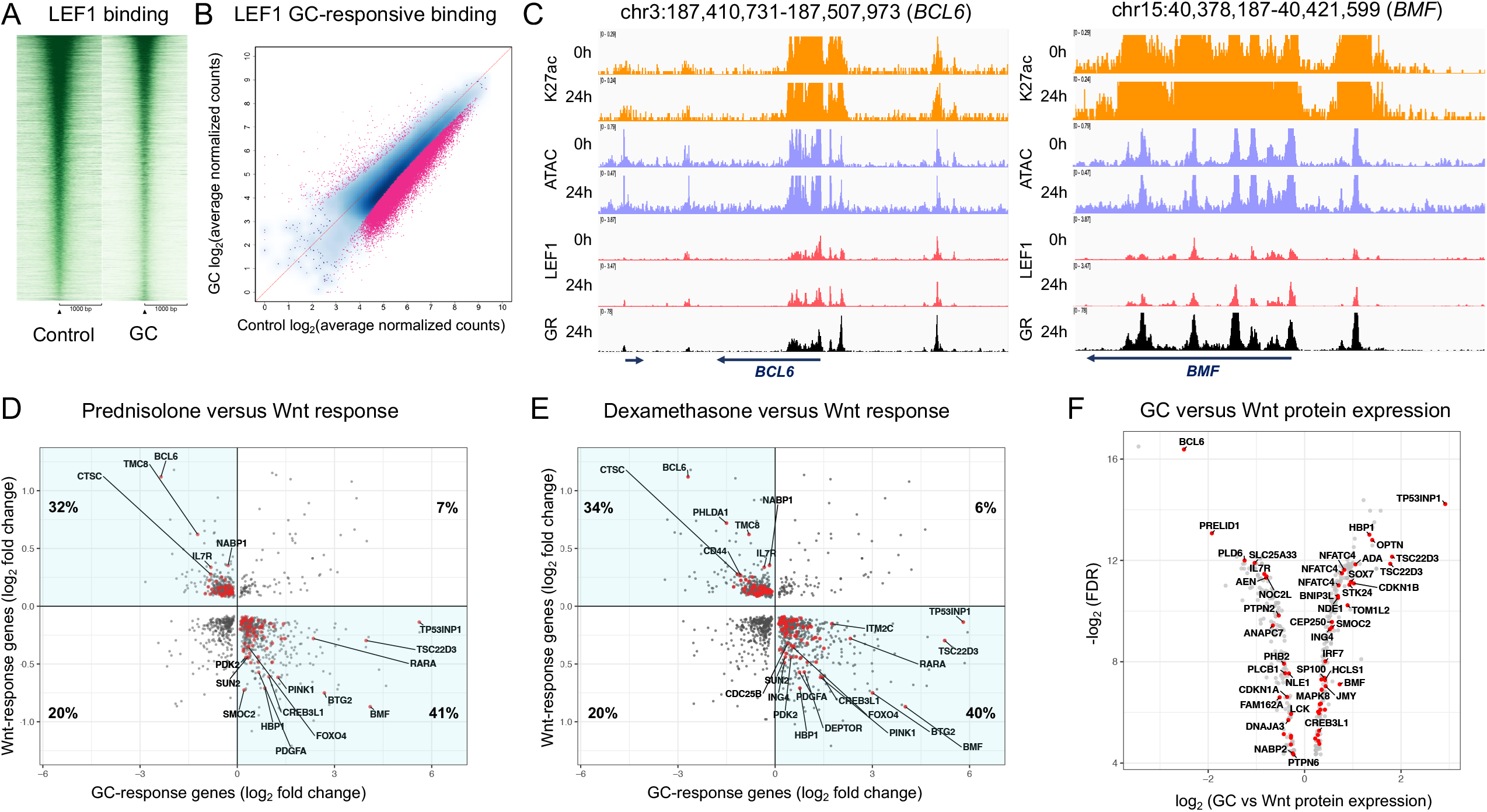
**(A)** CUT&RUN sequencing read enrichment at LEF1 binding sites +/- 1kb (Cell Signaling antibody # 2230). Enrichment is shown for a 24-hour treatment with vehicle control (left) or prednisolone (5uM; GC, right). **(B)** Log_2_-transformed normalized CUT&RUN sequencing read counts at LEF1 binding sites treated for 24 hours with vehicle control (x-axis) or prednisolone (y-axis). Binding sites exhibiting significant differences in occupancy (FDR<0.05) are shown in pink. **(C)** IGV genome browser images of signal tracks providing examples of GR and LEF1 co-occupancy at *BCL6* (left) and *BMF* (right) gene loci. **(D)** Log_2_-transformed fold changes of genes commonly regulated by GC and Wnt signaling pathways in Nalm6 cells after 24 hours of treatment using 5uM prednisolone (GC-response genes; x-axis) or 0.5uM CHIR-99021 (Wnt-response genes; y-axis). GC-response and Wnt-response gene log_2_ fold changes are provided. The percentage of genes in each quadrant is provided, and genes showing opposing effects are highlighted in quadrants 2 and 4. Red denotes genes involved in cell death, cell proliferation and/or cell cycle pathways and notable genes are labeled. **(E)** Log_2_-transformed fold changes of genes commonly regulated by GC and Wnt signaling pathways in Nalm6 cells after 24 hours of treatment using 100nM dexamethasone (GC-response genes; x-axis) or 0.5uM CHIR-99021 (Wnt-response genes; y-axis). GC-response and Wnt-response gene log_2_ fold changes are provided. The percentage of genes in each quadrant is provided, and genes showing opposing effects are highlighted in quadrants 2 and 4. Red denotes genes involved in cell death, cell proliferation and/or cell cycle pathways and notable genes are labeled. **(F)** Volcano plot of differential protein expression between prednisolone (5uM) or CHIR-99021 (0.5uM) treatment for 24 hours. Plot shows proteins of discordantly regulated prednisolone-response and Wnt-response genes with consistent directionality. Red denotes proteins involved in cell death, cell proliferation and/or cell cycle pathways and notable proteins are labeled.

To determine if overlapping TF occupancy impacts crosstalk between GC and canonical Wnt signaling transcriptional responses, RNA-seq was performed on WT Nalm6 cells treated for 24 hours with CHIR-99021 or vehicle control. We identified that 84% of Wnt-response genes were also prednisolone-response genes in Nalm6 cells. Supporting a mutual antagonism between these two signaling pathways, 73% of these common response genes exhibited discordant changes in gene expression (**Figure 2D**) and were enriched in cell death and cell cycle pathways (**supplemental Table 2**). Consistent patterns were also found using dexamethasone-response genes in Nalm6 cells (**Figure 2E**) and CEM T-ALL cells (**supplemental Figure 14**), with most Wnt-response genes overlapping GC-response genes, and most of these common response genes being discordantly regulated (Nalm6=74%, CEM=65%) and enriched for similar biological pathways (**supplemental Table 2**). Over 94% of discordantly regulated prednisolone-response genes were also discordantly regulated dexamethasone-response genes. Most discordantly regulated prednisolone-response genes (58%) were also associated with overlapping GR and LEF1 or GR and TCF7L2 binding events in Nalm6 cells. Sixty-one discordantly regulated prednisolone-response genes were also previously linked to GC resistance in primary B-ALL cells from over 200 patients [^7^; **supplemental Table 3**], supporting a broader impact on GC resistance, and most of these genes (75%; 46 of 61) were associated with overlapping GR and LEF1 or GR and TCF7L2 binding events. Most discordantly regulated GC-response genes (>=66%) also exhibited a more blunted GC response in TLE1 KO cells compared to WT cells, concordant with enhanced canonical Wnt signaling and reduced cellular apoptosis from TLE1 ablation (**supplemental Figure 15**). To determine if this transcriptional antagonism translates to the proteome level, we performed mass spectrometry in Nalm6 cells after a 24-hour treatment with prednisolone or CHIR-99021. We uncovered that 50% of discordantly regulated prednisolone-response genes also exhibited significantly different patterns as proteins (**supplemental Table 4**), and 66% of these proteins displayed consistent directionality (**Figure 2F**).

Collectively, our data uncovered extensive crosstalk and mutual antagonism between GC and canonical Wnt signaling pathways in B-ALL cells, and we confirmed similar effects in T-ALL cells. This antagonism is mediated in part through binding of GC and Wnt TFs to common *cis*-regulatory elements associated with cell death and cell proliferation genes. This overlap in TF occupancy was further accompanied by overlapping and opposing transcriptional programs that impacted protein expression. Overall, these data suggest *cis*-regulatory disruptions at *TLE1* are linked to GC resistance in primary B-ALL cells from patients through reduced GC-mediated apoptosis via enhanced canonical Wnt signaling. As a result of the deep genomic and gene regulatory connectivity between these two signaling pathways, our data supports the importance of canonical Wnt signaling in mediating GC resistance in B-ALL, and further suggest that treatment with canonical Wnt antagonists may improve GC sensitivity in patients with resistant disease.

## Disclosures

No conflicts of interest to disclose.

## Contributions

Conceptualization, D.S.; Methodology, B.P.B, K.R. Barnett, K.R. Bhattarai, R.J.M., A.A.H., J.P., S.M.P., D.S.; Investigation, B.P.B, K.R. Barnett, K.R. Bhattarai, R.J.M., B.S.H., A.B., K.K.; Formal Analysis, B.P.B, K.R. Barnett, K.R. Bhattarai, R.J.M., A.B., K.K., A.A.H., D.S.; Clinical data acquisition, S.J., C.H.P; Writing – Original Draft, B.P.B, D.S.; Writing – Review & Editing, B.P.B, K.R. Barnett, K.R. Bhattarai, R.J.M., B.S.H., A.B., K.K., A.A.H., S.J., C.H.P, J.P., S.M.P., D.S.; Funding Acquisition, D.S.

## Data-sharing statement

All functional genomic data from cell lines have been deposited into the Gene Expression Omnibus (GSE215188).

## Acknowledgements

We would like to thank the St. Jude Hartwell Center for next-generation sequencing, the St. Jude Center for Advanced Genome Engineering for CRISPR/Cas9 genome editing and the St. Jude Center for Proteomics and Metabolomics for mass spectrometry. This work was supported by the National Institutes of Health (R01CA234490 and P30CA021765) and the American Lebanese Syrian Associated Charities (ALSAC). The content of this paper is solely the responsibility of the authors and does not necessarily represent the official views of the U.S. National Institutes of Health.

## Notes

### Competing Interest Statement

The authors have declared no competing interest.

### Summary of Updates

A revised version of the manuscript has been uploaded which includes new analyses and experimentation. Figures 1 and 2 have been revised. Eleven additional supplemental figures and three additional supplemental tables have now also been included.

