## Supplemental Figures for "Mutual antagonism between glucocorticoid and canonical Wnt signaling pathways in B-cell acute lymphoblastic leukemia"

[1] Hematological Malignancies Program, St. Jude Children's Research Hospital, Memphis, TN 38105, USA.
[2]Department of Pharmacy and Pharmaceutical Sciences, St. Jude Children's Research Hospital, Memphis, TN 38105, USA.
[3]Graduate School of Biomedical Sciences, St. Jude Children's Research Hospital, Memphis, TN 38105, USA.
[4]Center for Advanced Genome Engineering, St. Jude Children's Research Hospital, Memphis, TN 38105, USA.
[5]Center for Proteomics and Metabolomics, St. Jude Children's Research Hospital, Memphis, TN 38105, USA.
[6]Department of Oncology, St. Jude Children's Research Hospital, Memphis, TN 38105, USA.
[7]Department of Structural Biology, St. Jude Children's Research Hospital, Memphis, TN 38105, USA.
[8]Department of Developmental Neurobiology, St. Jude Children's Research Hospital, Memphis, TN 38105, USA.
[9]Department of Cell and Molecular Biology, St. Jude Children's Research Hospital, Memphis, TN 38105, USA.
[10]Integrated Biomedical Sciences Program, University of Tennessee Health Science Center, Memphis, TN 38105, USA.
[*]Corresponding author

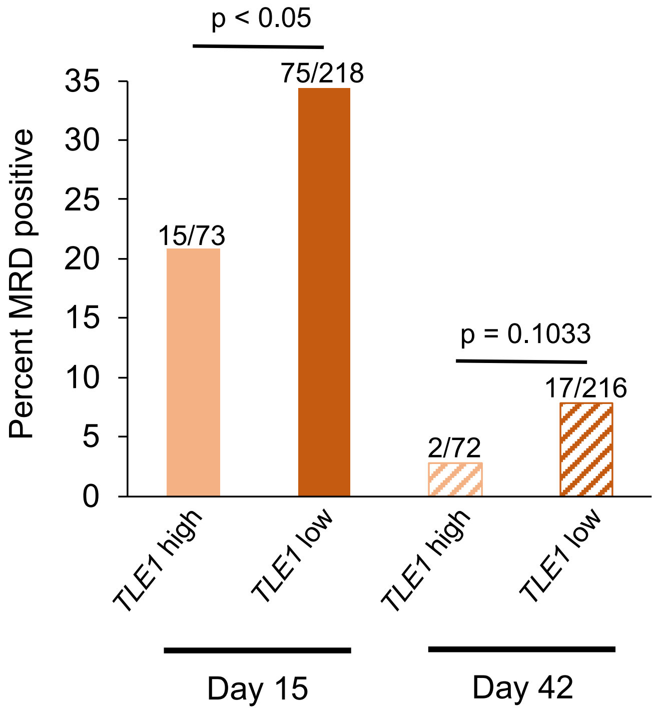


**Supplemental Figure 1.** Barplot showing the percent of minimal residual disease (MRD) positive patients at day 15 (left; MRD ≥ 1%) or day 42 (right; MRD ≥ 0.1%) with primary leukemic cells harboring high versus low baseline *TLE1* expression. RNA-seq derived *TLE1* expression above the 3rd quartile was calculated using RNA-seq data across all patient leukemic cells and was used as a cutoff for high *TLE1* expression. P-values are provided that test the hypothesis that higher *TLE1* expression is associated with reduced MRD positive cases (Fisher's Exact, one-sided).

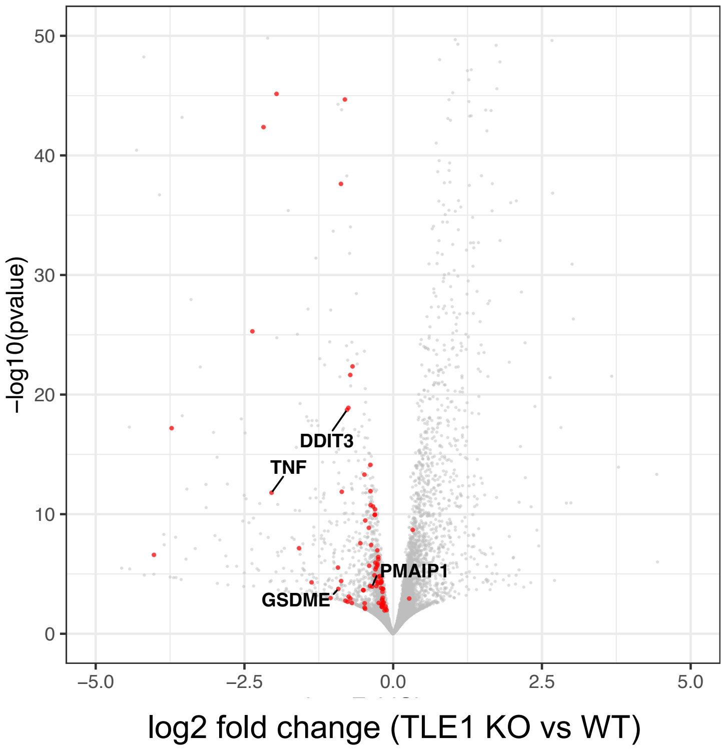


**Supplemental Figure 2.** Volcano plot showing differentially expressed genes between Nalm6 WT and TLE1 KO cells in the absence of GCs. Genes involved in apoptotic pathways are shown in red and several notable genes are highlighted.

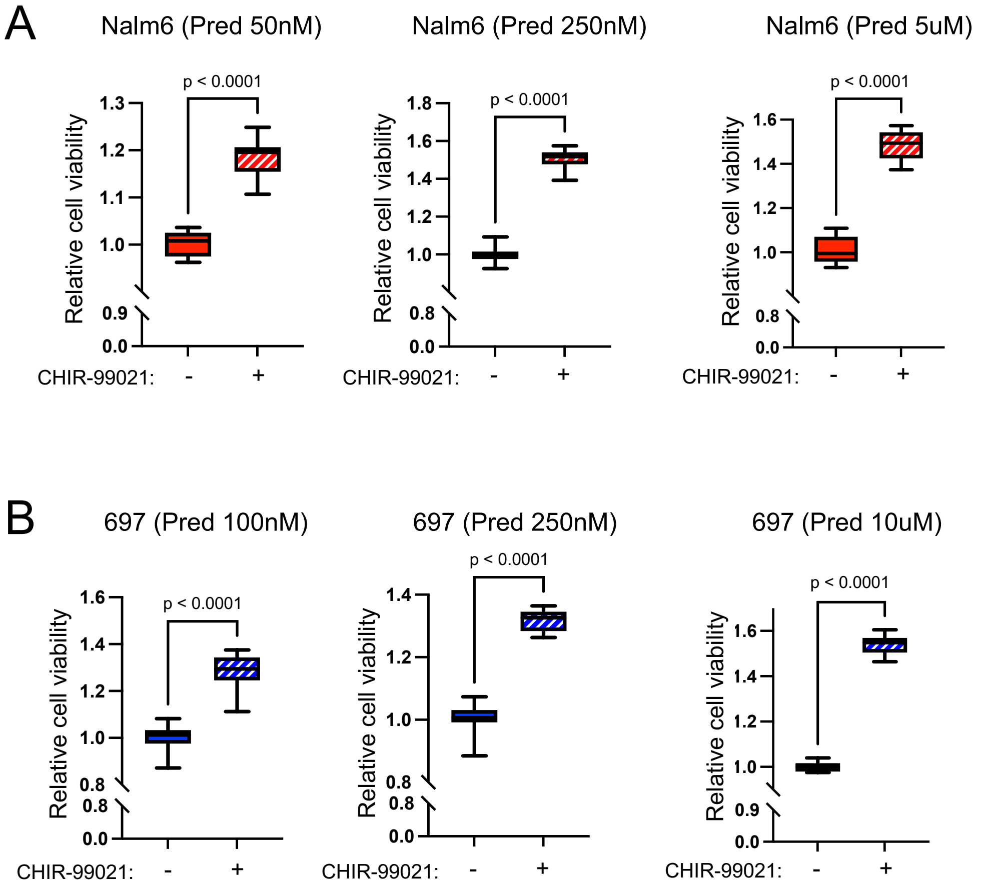


**Supplemental Figure 3. (A)** GC drug viability results displaying the relative viability for Nalm6 cells treated with only prednisolone (50nM, 250nM, or 5uM) or in combination with the canonical Wnt agonist CHIR-99021 (0.5uM) after 72-hours of treatment, n = 12 per group. **(B)** GC drug viability results displaying the relative viability for 697 cells treated with only prednisolone (100nM, 250nM, or 10uM) or in combination with the canonical Wnt agonist CHIR-99021 (0.5uM) after 72-hours of treatment, n = 12 per group.

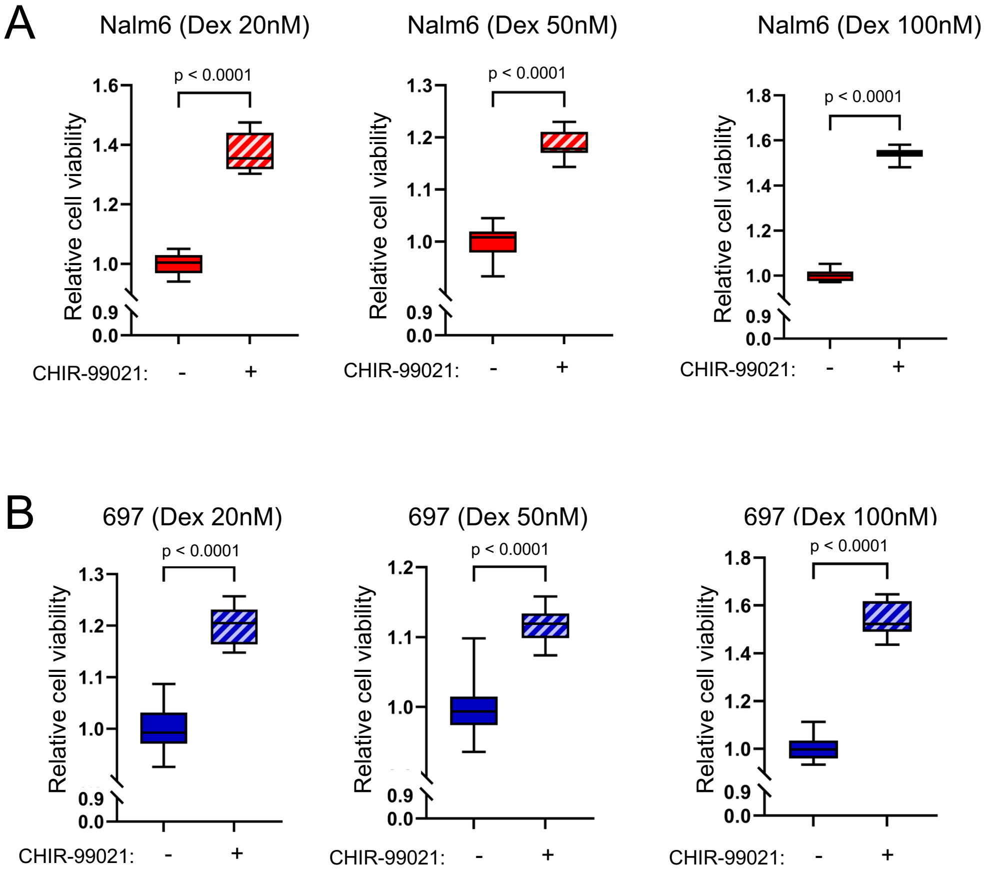


**Supplemental Figure 4. (A)** GC drug viability results displaying the relative viability for Nalm6 cells treated with only dexamethasone (20nM, 50nM, or 100nM) or in combination with the canonical Wnt agonist CHIR-99021 (0.5uM) after 72-hours of treatment, n = 12 per group. **(B)** GC drug viability results displaying the relative viability for 697 cells treated with only dexamethasone (20nM, 50nM, or 100nM) or in combination with the canonical Wnt agonist CHIR-99021 (0.5uM) after 72-hours of treatment, n = 15 per group.

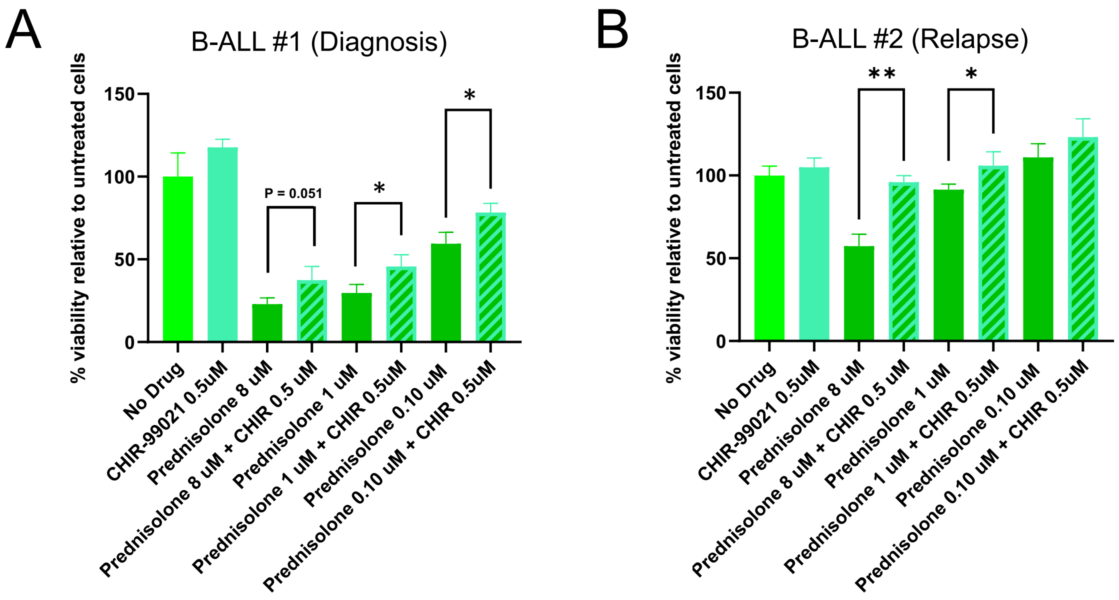


**Supplemental Figure 5.** GC drug viability results displaying the percent viability relative to untreated control cells for distinct patient-derived xenograft primary B-ALL cells harvested from two patients at diagnosis **(A)** or after relapse **(B).** Cells were treated *ex vivo* with only prednisolone (0.1uM, 1uM, or 8uM) or in combination with the canonical Wnt agonist CHIR-99021 (0.5uM) for 72-hours, n = 3 per group. * = p<0.05, ** = p<0.01.

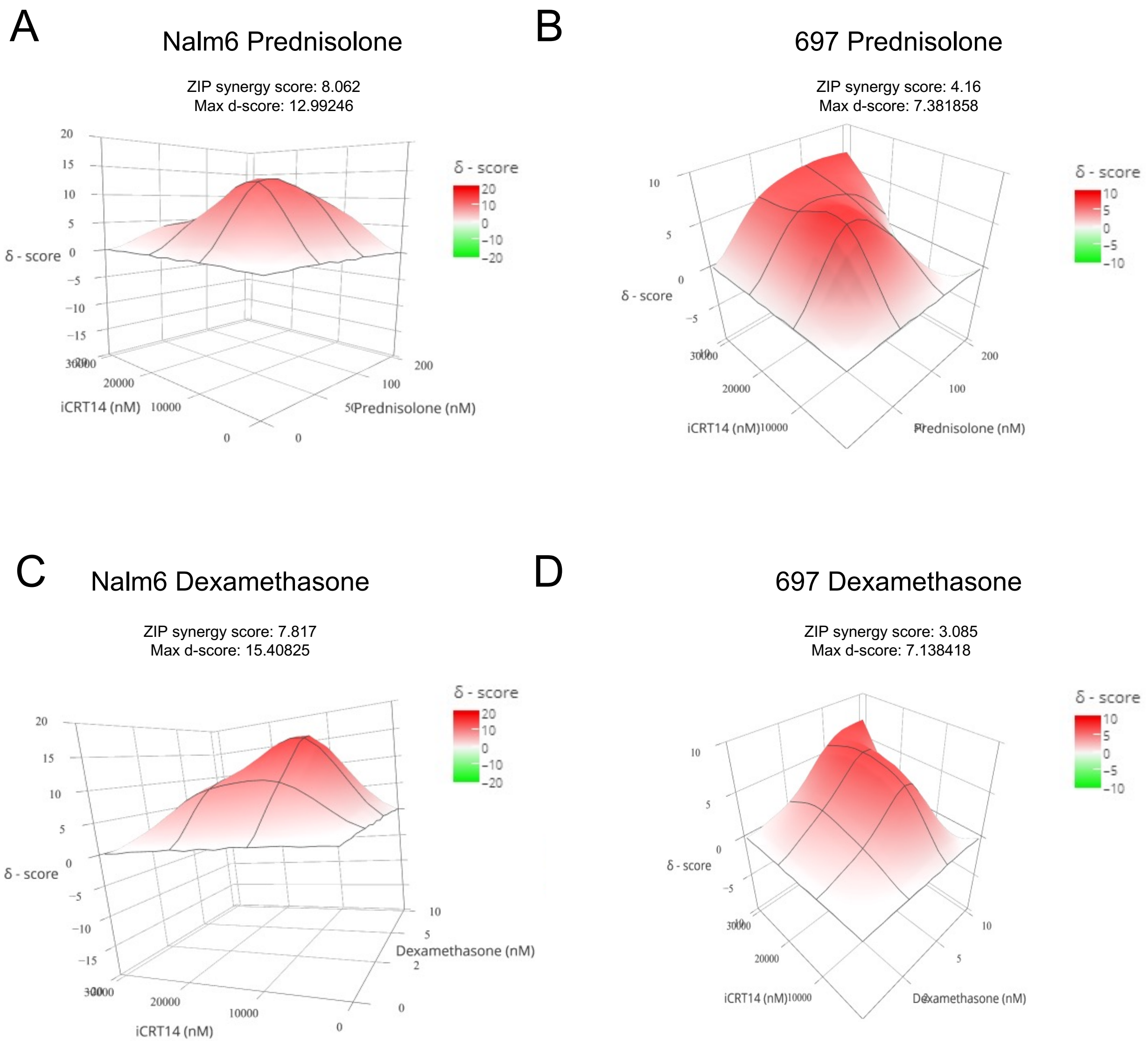


**Supplemental Figure 6.** Analysis of the effects of drug co-treatment in Nalm6 **(A,C)** and 697 **(B,D)** human B-ALL cell lines using prednisolone **(A,B)** or dexamethasone **(C,D)** and the canonical Wnt antagonist iCRT14 for 72 hours, n = 3 per group. Concentrations of glucocorticoid and iCRT14 used in co-treatment are provided on x and z-axes, respectively. Delta scores (d-score or δ - score) are provided on the Y-axis. The maximum d-score and Zero Interaction Potency (ZIP) score are provided above each graph. Analysis was performed by SynergyFinder web application: http://synergyfinder.fimm.fi/.

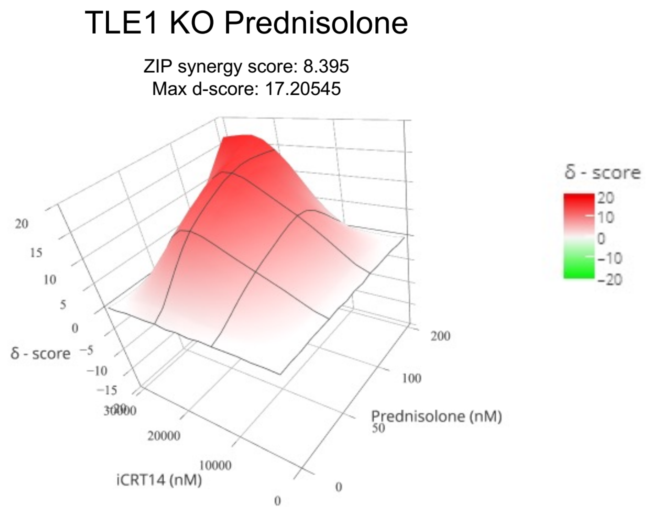


**Supplemental Figure 7.** Analysis of the effects of drug co-treatment in TLE1 KO Nalm6 cell lines using prednisolone and the canonical Wnt antagonist iCRT14 for 72 hours, n = 3 per group. Concentrations of prednisolone and iCRT14 used in co-treatment are provided on x and z-axes, respectively. Delta scores (d-score or δ - score) are provided on the Y-axis. The maximum d-score and Zero Interaction Potency (ZIP) score are provided above each graph. Analysis was performed by SynergyFinder web application: http://synergyfinder.fimm.fi/.

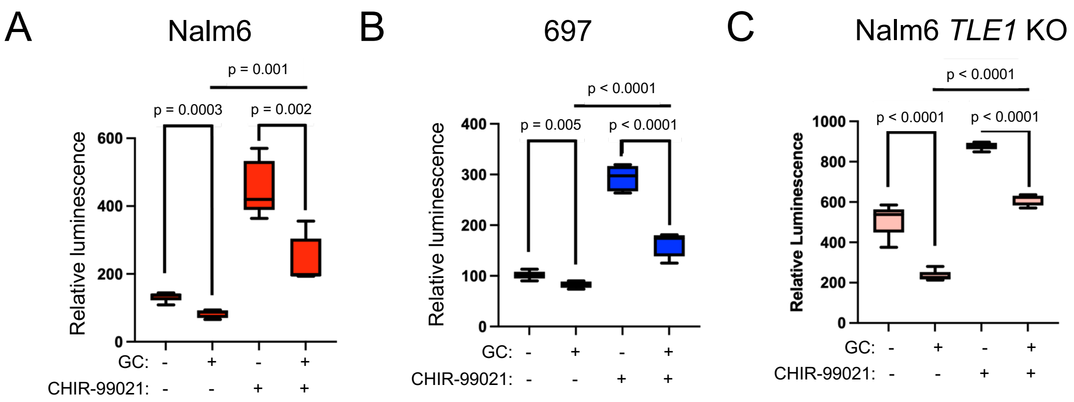


**Supplemental Figure 8.** **(A)** beta-catenin luciferase reporter assay for Nalm6 cells treated with prednisolone (GC, 5uM), CHIR-99021 (0.5uM), or both for 24 hours, n = 5 per group. **(B)** Beta-catenin luciferase reporter assay for 697 cells treated with prednisolone (GC, 10uM), CHIR-99021 (0.5uM), or both for 24 hours, n = 5 per group. **(C)** beta-catenin luciferase reporter assay for TLE1 *KO* Nalm6 cells treated with prednisolone (GC, 5uM), CHIR-99021 (0.5uM), or both for 24 hours, n=5 per group.

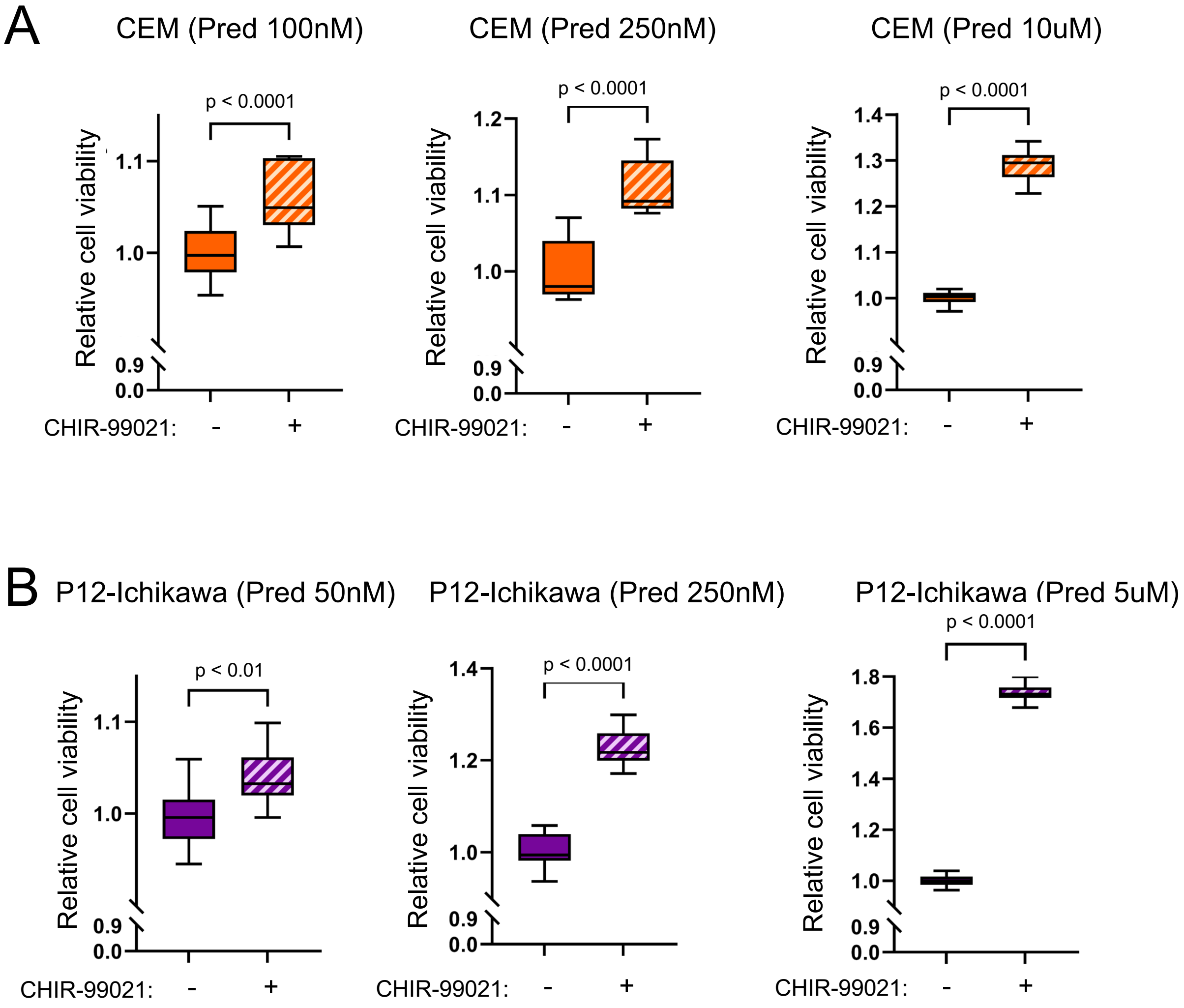


**Supplemental Figure 9. (A)** GC drug viability results displaying the relative viability for CEM T-ALL cells treated with only prednisolone (100nM, 250nM, or 10uM) or in combination with the canonical Wnt agonist CHIR-99021 (2uM) after 72 hours of treatment, n = 15 per group. **(B)** GC drug viability results displaying the relative viability for P12-Ichikawa T-ALL cells treated with only prednisolone (50nM, 250nM, or 5uM) or in combination with the canonical Wnt agonist CHIR-99021 (10uM) after 72 hours of treatment, n = 15 per group.

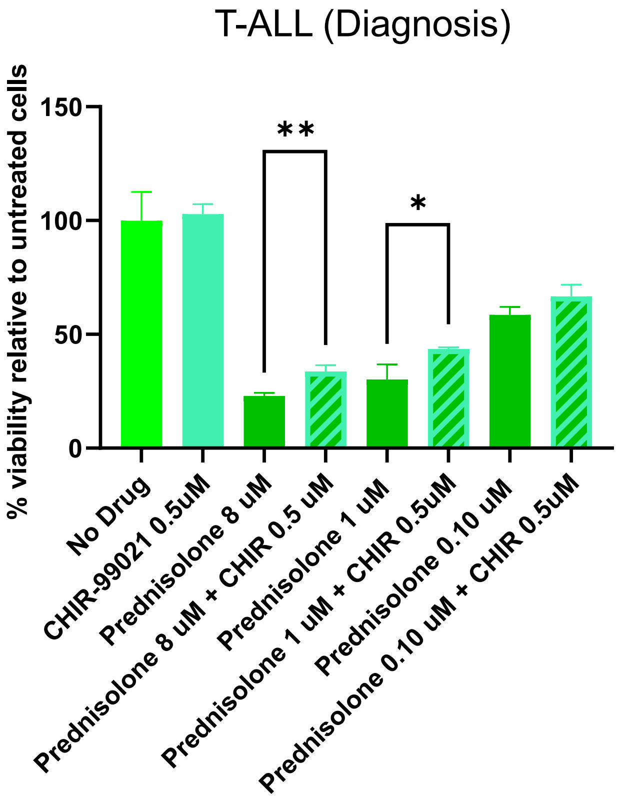


**Supplemental Figure 10.** GC drug viability results displaying the percent viability relative to untreated control cells for patient-derived xenograft primary T-ALL cells harvested from patients at diagnosis**.** Cells were treated *ex vivo* with only prednisolone (0.1uM, 1uM, or 8uM) or in combination with the canonical Wnt agonist CHIR-99021 (0.5uM) for 72-hours, n = 3 per group. * = p<0.05, ** = p<0.01.

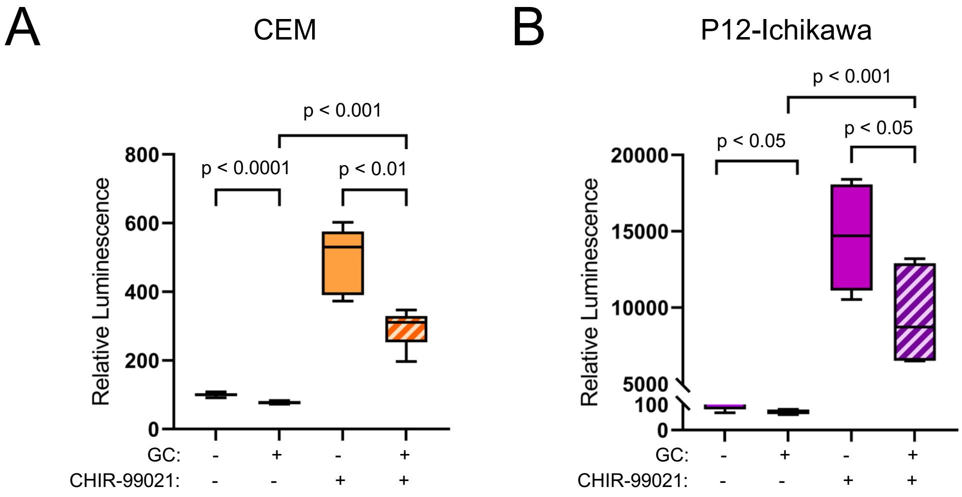


**Supplemental Figure 11. (A)** Beta-catenin luciferase reporter assay for CEM cells treated with prednisolone (GC, 10uM), CHIR-99021 (2uM), or both for 24 hours, n = 6 per group. **(B)** Beta-catenin luciferase reporter assay for P12-Ichikawa cells treated with prednisolone (GC, 5uM), CHIR-99021 (10uM), or both for 24 hours, n = 6 per group.

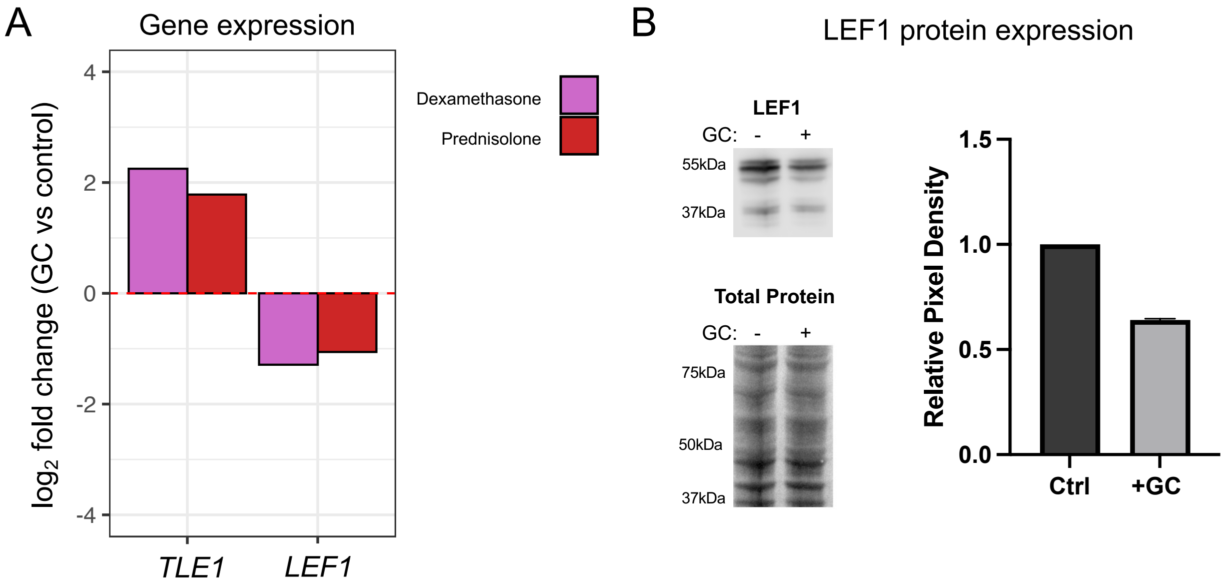


**Supplemental Figure 12. (A)** $Log_2$ fold changes of significant differential expression (RNA-seq FDR<0.05) for *TLE1* and *LEF1* in Nalm6 cells after 24 hours of prednisolone (5uM; red) or dexamethasone (100nM; purple) treatment. **(B)** Protein expression of LEF1 (Cell Signaling antibody # 2230). Western blot on wild-type cell lysates treated with vehicle control (-) or prednisolone (5uM; +) for 24 hours is shown at the top and image of total protein is shown below. Quantitation blot image for LEF1 is provided on the right. Standard deviation error bars are shown, n=2 per group.

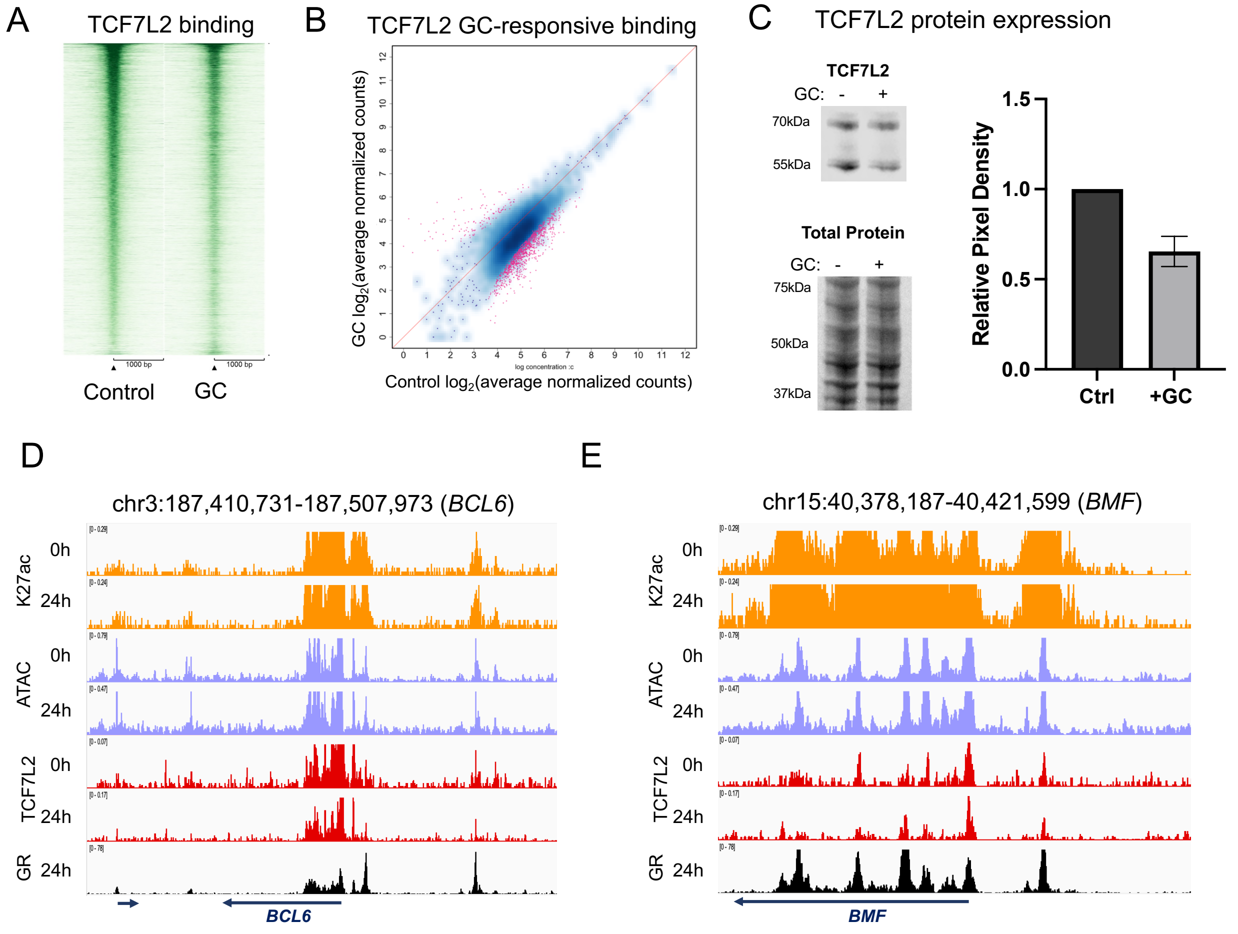


**Supplemental Figure 13. (A)** CUT&RUN sequencing read enrichment at TCF7L2 binding sites +/- 1kb (Cell Signaling antibody # 2230). Enrichment is shown for a 24-hour treatment with vehicle control (left) or prednisolone (5uM; GC, right). **(B)** $Log_2$-transformed normalized CUT&RUN sequencing read counts at TCF7L2 binding sites treated for 24 hours with vehicle control (x-axis) or prednisolone (y-axis). Binding sites exhibiting significant differences in occupancy (FDR<0.05) are shown in pink. **(C)** Protein expression of TCF7L2 (Cell Signaling antibody # 2569). Western blot on wild-type cell lysates treated with vehicle control (-) or prednisolone (5uM; +) for 24 hours is shown at the left. Western blot on wild-type cell lysates treated with vehicle control (-) or prednisolone (5uM; +) for 24 hours is shown at the top and image of total protein is shown below. Quantitation of blot image for TCF7L2 is provided on the right. Standard deviation error bars are shown, n=2 per group. IGV genome browser images of signal tracks providing examples of GR and TCF7L2 co-occupancy at *BCL6* **(D)** and *BMF* **(E)** gene loci.

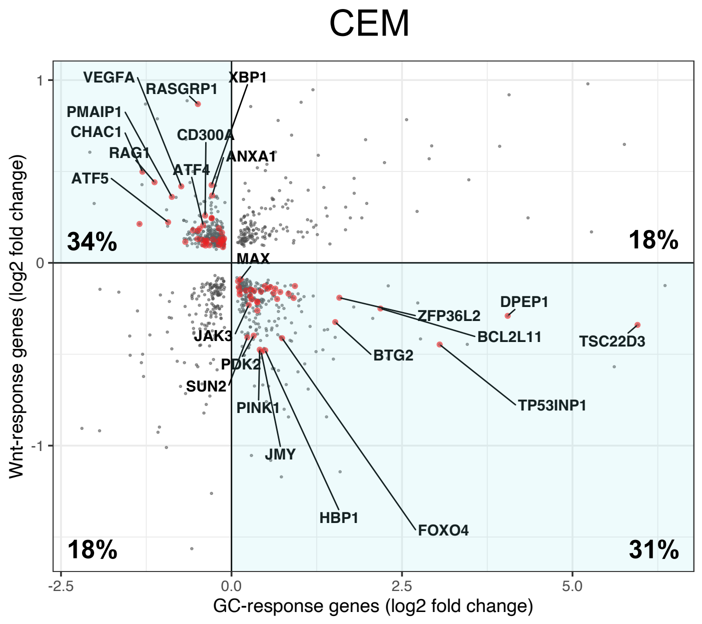


**Supplemental Figure 14.** $Log_2$-transformed fold changes of genes commonly regulated by GC and Wnt signaling pathways in CEM cells after 24 hours of treatment using 10uM prednisolone (GC-response genes; x-axis) or 2uM CHIR-99021 (Wnt-response genes; y-axis). GC-response and Wnt-response gene $log_2$ fold changes are provided. The percentage of genes in each quadrant is provided, and genes showing opposing effects are highlighted in quadrants 2 and 4. Red denotes genes involved in cell death, cell proliferation and/or cell cycle pathways and notable genes are labeled.

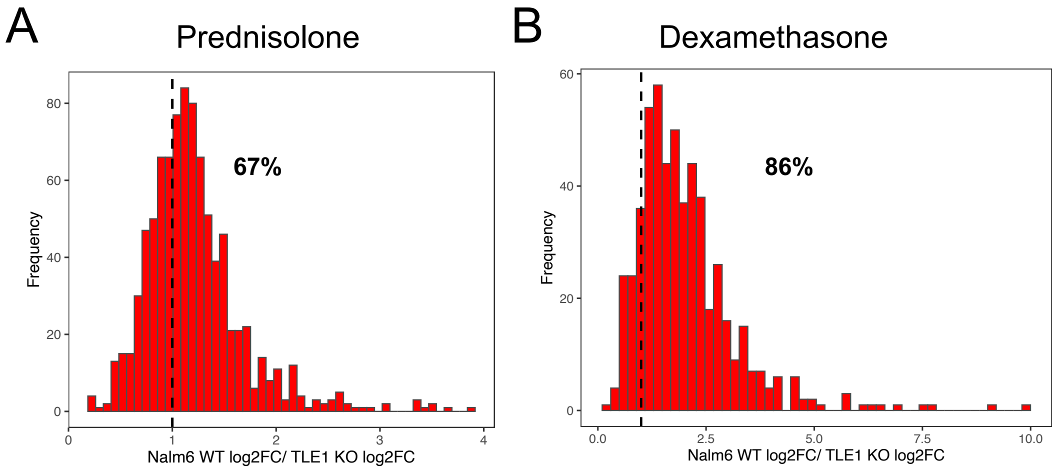


**Supplemental Figure 15.** Comparison of $Log_2$-transformed fold changes of discordantly regulated genes between Nalm6 WT and Nalm6 TLE1 KO cells after 24 hours of treatment using 5uM prednisolone (**A**) or 100nM dexamethasone (**B**). The percentage of discordantly regulated genes with greater fold change in WT cells compared to TLE1 KO cells is provided.
